# The Rise of the Middle Author: Investigating Collaboration and Division of Labor in Biomedical Research using Partial Alphabetical Authorship

**DOI:** 10.1101/108357

**Authors:** Philippe Mongeon, Elise Smith, Bruno Joyal, Vincent Larivière

## Abstract

Contemporary biomedical research is performed by increasingly large teams. As a consequence, an increasingly large number of individuals are being listed as authors in the byline of biomedical articles, which complicates the proper attribution of credit and responsibility to individual authors for their work. Typically, more importance is given to the first and last authors of biomedical papers, and the others (the middle authors) are considered to have made smaller contributions. However, we argue that this distinction between first, middle and last authors does not properly reflect the actual division of labor and does not allow a fair allocation of credit among the members of the research teams. In this paper, we use partial alphabetical authorship to divide the authors of all biomedical articles in the Web of Science published over the 1980-2015 period in three groups: primary authors, middle authors, and supervisory authors. We show that alphabetical ordering of middle authors is frequent in biomedical research, and that the prevalence of this practice is positively correlated with the number of authors in the bylines. We also find that, for articles with 7 or more authors, the average proportion of team members in each group is independent of the team size, more than half of the authors being middle authors. This suggests that growth in authors’ lists are not due to an increase in secondary contributions but, rather, in equivalent increases of all types of roles and contributions. Nevertheless, we show that the relative contribution of middle authors to the overall production of knowledge in the biomedical field as increased dramatically over the last 35 years.

## Introduction

With the increasing costs, complexity and interdisciplinarity of modern science [1], research collaboration has become the norm [2]. Scientific knowledge is now being produced by increasingly large teams [3,4], generally involving researchers from multiple disciplines, institutions and countries [5]. Funding agencies often encourage and facilitate collaboration [6–8] and there is evidence that funded research is indeed more collaborative [9,10]. A growing body of evidence also suggests that collaborative research has more impact and that increasingly large and diverse teams are necessary to achieve greater impact [4].

Increased collaboration and larger teams translate into a larger number of authors listed in the byline of scholarly articles. In certain cases, there may be hundreds of authors on a paper; a phenomenon coined as ‘hyperauthorship’ [11]. This alone, but also the diversity of collaboration types [12], the team composition [13], and the division of the work within the team [14], greatly complicates the attribution of credit and responsibility among the team members [15]. This is an important issue since the advancement of researchers’ careers largely depends on the credit they obtain for their work [16,17]. Because authorship is so important, conflicts regarding authorship are becoming commonplace [16,17] and may introduce a tension within the workplace. The growing complexity of credit attribution is also potentially detrimental for the scientific system as a whole, which works best when excellence is properly identified and rewarded [18].

While it may be difficult for an external observer to assess with precision the respective contribution of individual authors of a collaborative work, there are implicit disciplinary norms regarding authorship of individual authors on the byline as a proxy for the extent and nature of their respective contributions is possible because names are typically ordered following implicit disciplinary norms [19]. For example, in the biomedical field, as in most lab-based disciplines, authorship order is based on the importance and type of the contribution as well as the hierarchical position within the team or laboratory. Generally, the first and last position are given the most importance. The first author is a PhD student or a postdoctoral fellow who contributed most to the research, and the last is generally the lab director [20]. Between the first and last authors are listed an increasingly large number of ‘middle authors’ who have typically played a less significant role in the research [14]. A number of scientific journals (e.g., JAMA, BMJ, the Lancet, NEJM and PLoS) require author’s contribution statement intended to provide useful information about an individual author’s contribution. However, the value of contribution statements is limited because of significant reporting biases (Ivaniš, Hren, Sambunjak, Marušić & Marušić, 2008; Ivaniš, Hren, Marušić & Marušić, 2011), and because the statements address the type of work performed but not on the relative value or importance of the work. Nonetheless, several analyses of the relation between the authors rank on the byline and their reported contributions [14,e.g.,21] confirmed the polarization of ‘core’ contributors towards the first and the last position of the authors list, while authors who made fewer types of contributions were listed in the middle. In this paper, we divide the bylines of biomedical articles into three distinct groups using a terminology similar to the one proposed by Baerlocher and colleagues [21]:

1) *Primary authors*: main contributors to the experimental work;
2) *Supervisory authors*: senior researchers who supervised the research; and
3) *Middle authors*: individuals with relatively small contributions to the research who are listed between the primary and supervisory authors.

While such a division of the bylines might better reflect the work performed by the authors, and allow a more adequate and fair attribution of credit, it also raises a difficult question: how can we distinguish primary, middle and supervisory authors? Previous bibliometric analyses of biomedical research [22,e.g.,23,24] have typically divided the bylines in three parts: the first author, the middle authors, and the last author. This poorly reflects reality since it allows only be one primary author and only one supervisory author. This is problematic as collaborative research (especially inter-institutional or interdisciplinary research) is likely to have multiple primary authors leading perhaps different part of the experimental work, and also multiple supervisory authors [25]. Thus, the ‘first author + middle authors + last author’ model is an arbitrary division of authors that might unfairly tag as middle authors some researchers who played major roles in the research.

In this paper, we use partial alphabetical authorship as a tool to delineate primary, middle and supervisory authors. As Harriet Zuckerman [26] pointed out, listing a subset of authors in alphabetical order creates a clear distinction between those who are listed alphabetically and those who are not. For instance, if an article has twenty authors, and the six main contributors (the first four and the last two) are not listed in alphabetical order, while authors from the fifth to the eighteenth position are, a distinction is made; identifying sequences of authors in alphabetical order in the middle of the bylines serves to distinguish the primary, middle and supervisory authors. This method assumes that the contributions of authors listed in alphabetical order were not as substantial as that of the others.

The identification of primary middle and supervisory authors through the use of alphabetical subsets does, however, have certain limitations. One might argue that there are other reasons for this type of ordering, such as the existence of large teams in which many authors have contributed almost equally to different parts of a project. While ordering the first few authors according to their contributions might seem like a necessary burden, researchers may at some point decide that it is simpler – and maybe fairer – to list a subset of authors alphabetically, thus differentiating these authors from others.

The purpose of this study is to empirically explore the relative contribution of primary authors, middle authors and supervisory authors in the biomedical field. More specifically we provide answers to the following research questions:

1. How prevalent are alphabetically ordered middle authors in biomedical research?
2. What are the proportions of primary, middle and supervisory authors in the articles’ bylines?
3. What is the overall contribution of middle authors to the biomedical literature?

## Methods

### Data

This study is based on all biomedical research and clinical medicine articles published between 1980 and 2015, which were authored by 4 to 100 individuals, and indexed in Thomson Reuters’ Web of Science (WoS). The discipline of the articles was determined by the NSF classification of the journal in which they are published. Because trends observed were almost identical in the two biomedical disciplines studied (Biomedical Research and Clinical Medicine), the results presented below are for the two disciplines combined. We identified middle authors using the following these three steps: 1) identify alphabetical sequences, 2) correct broken sequences, and 3) distinguish intentional and incidental alphabetical sequences.

### Identifying Middle Authors

We used an approach similar to that of Waltman [27] to detect sequences of authors in alphabetical order by giving each author of a byline an alphabetical rank based on their last name, and then their initials. An alphabetically ordered sequence of authors is formed when a list of consecutively in alphabetically order. Consider for example, an article authored by Wilson, B., Smith, J., Albert, S., Carter, B., Miller, D., Ford, R., and Clark, P.; it includes a grouping of three authors (Albert, S., Carter, B., and Miller, D.) in alphabetical order starting from the 3^rd^ position and ending at the 5^th^ position of the list.

### Correcting Broken Sequences

Depending solely on names and initials to identify alphabetical sequences has some limitations. Errors can occur because of special character conversion, compound names and names with prefixes, indexation errors and human errors in the alphabetical ordering. In our dataset, spaces and hyphens are removed from last names (e.g. van Gogh becomes vanGogh), and special characters are converted into the basic Latin alphabet (e.g. Lübeck becomes Luebeck). Also, the prefixes of Dutch names (e.g., van, von, etc.) are not taken into account in the alphabetical ordering. It may also happen that the first of two last names of an author is treated as a second first name during the indexation process. Fig 1 shows an example where authors from the second to the second to last positions have been ordered alphabetically. However, the sequence breaks at the 10^th^ author (Starr Koslow Mautner) because her last name (Koslow) has been indexed as a second initial. There may also be cases of human errors, for example when two names are inverted in a long list of otherwise alphabetically ordered authors. Alphabetical ordering conventions differ by language and country, so different individuals may alphabetically order the same list of names in a different way. These conventions also contain rules regarding alphabetical ordering of special characters, which can create further errors since these characters are no longer present in the indexed names.

**Fig 1.**
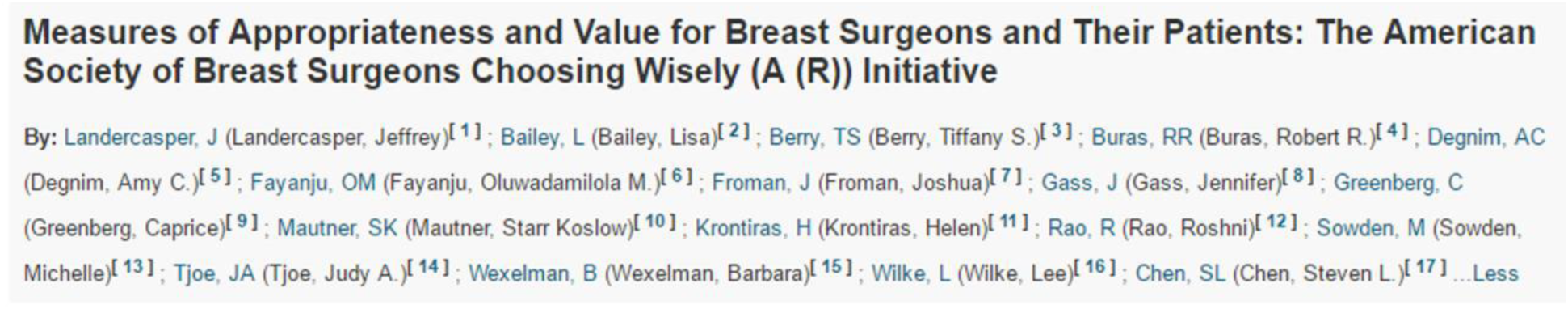
Example of a sequence break due to multiple last names.

To avoid, to the extent possible, the occurrence of the errors mentioned above we concatenated consecutive alphabetically ordered sequences which met one of the following conditions:

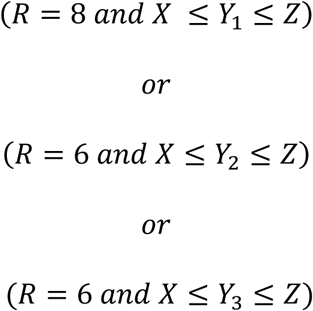

Where:

- *R* is the combined length (*r*) of the alphabetical sequences preceding and following the break.
- *X* is the first letter of the author name before the one causing the break.
- *Y1* is the first letter of the author name causing the break.
- *Y2* is the first letter of the author name causing the break after removing potential prefixes.
- *Y3* is the last initial of the author name causing the break.
- *Z* is the first letter of the author name after the one causing the break.

The value of *R* is important because the longer the consecutive sequences, the higher the probability that they actually constitute a single sequence that has been broken into two distinct parts. Therefore, to maximize the precision of the alphabetical sequence break detection method, we manually verified a random sample of 100 broken alphabetical sequences for different values of *R*, and we selected the minimum value of *R* for which the proportion of false positive was 5% or lower. A total of 192,716 broken alphabetical sequences were fixed: 28,779, 77,332 and 86,605 sequences with the (*R* = 8 *and X* ≤ *Y*_1_ ≤ *Z*), (*R* = 6 *and X* ≤ *Y*_2_ ≤ *Z*), and (*R* = 6 *and X* ≤ *Y*_3_ ≤ *Z*) conditions, respectively. The resulting dataset comprises more than 6.7 million articles authored by a total of more than 44 million authors, among which 13 million alphabetical sequences where found.

### Probability of Intentional vs. Chance Alphabetical Order

There is always a possibility that a list of authors in alphabetical order, which we use as a sequence of ‘middle authors’, is actually a pure result of chance and not intentional. Distinguishing intentional and chance alphabetical order is crucial since alphabetical sequences that occur randomly cannot be used to distinguish middle authors from the others. Thus, for each sequence found, we calculated *Pi*, which is the probability that the authors are intentionally listed in alphabetical order, and the opposite of the probability *Pc* that authors are listed alphabetically by chance (see S1 for details on the calculation of *Pi* and *Pc*). *Pi* is determined by two variable: the number of authors in the sequence (*r*) and the total number of authors in the byline (*N*), hereafter referred to as the team’s size. For example, there are 3,628,800 possible combinations of *N* = 10 authors out of which 156,002 contain an alphabetical sequence of *r* = 5 authors. Thus, a sequence of *r* = 5 has a 156,002/3,628,800 = 4.3% probability of occurring by chance (*Pc*), and therefore a 95.7% chance of being intentional (*Pi*). Fig 2 shows the relation between *N* and *Pi* for different values of *r*. We see that for short alphabetical sequences of 3 or 4 authors *Pi* increases rapidly as the byline gets longer, while the *Pi* of sequences of 6 and 7 remains very high, even for articles with up to 100 authors.

**Fig 2.**
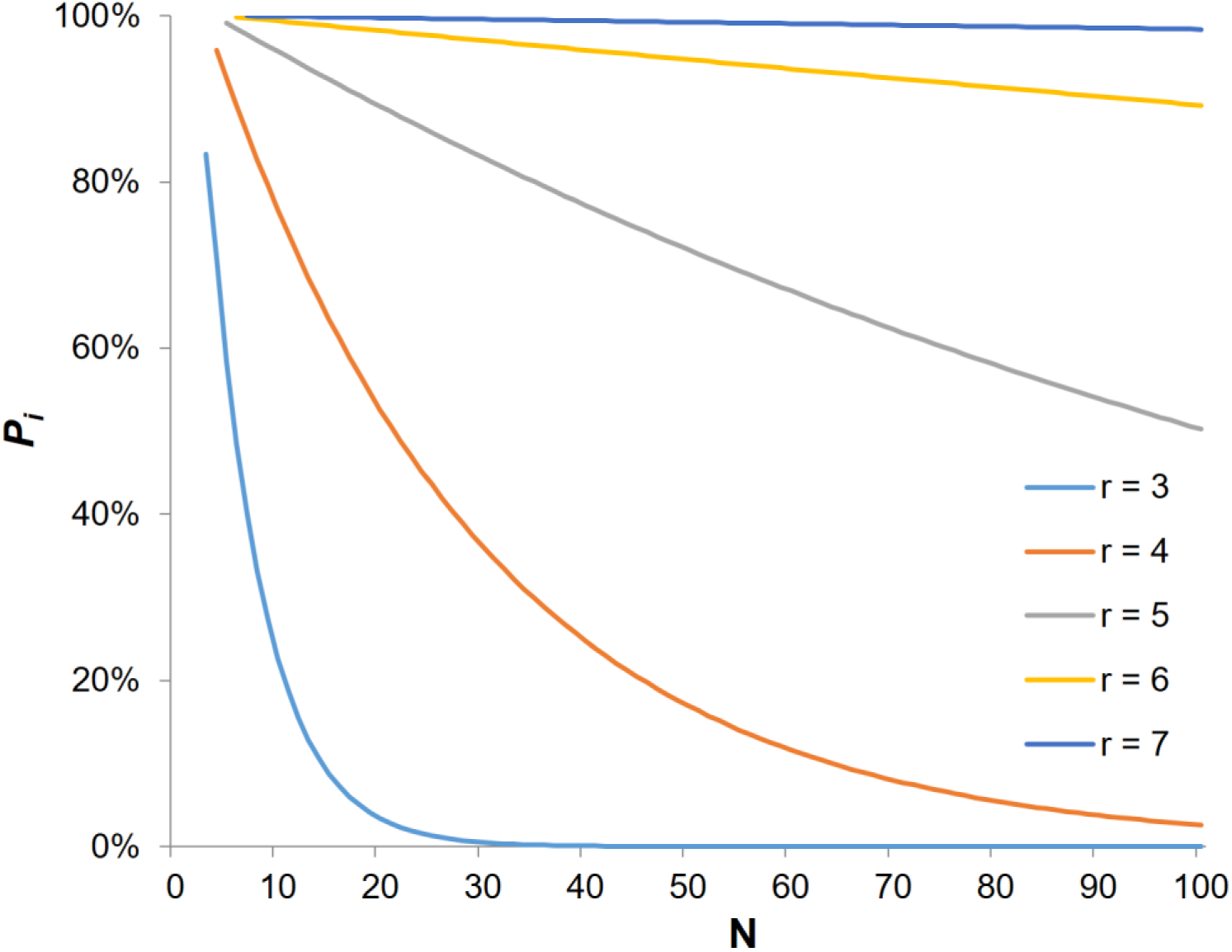
Probability of intentional alphabetical order (*P_i_*) as a function of the number of authors in the byline (N) and the length of the alphabetical sequence (r).

## Results

Fig 3 shows the probability of finding an intentional alphabetical sequence in articles’ bylines as a function of the team size (left) and as a function of the publication year (right). We distinguish here cases where middle authors are listed in alphabetical order from cases where the alphabetical sequence begins with the first author or ends with the last authors, as well as cases where all authors are in alphabetical order. Results suggest that alphabetical sequences occur more frequently in the middle of the authors list, and that their prevalence correlates with the team size. The average *Pi* quickly reaches 25% at *N* = 7 and increases to 50% for *N* = 35. This confirms that it is indeed common practice, in the biomedical field, to order middle authors in alphabetical order. The other types of alphabetical patterns remain relatively rare.

**Fig 3.**
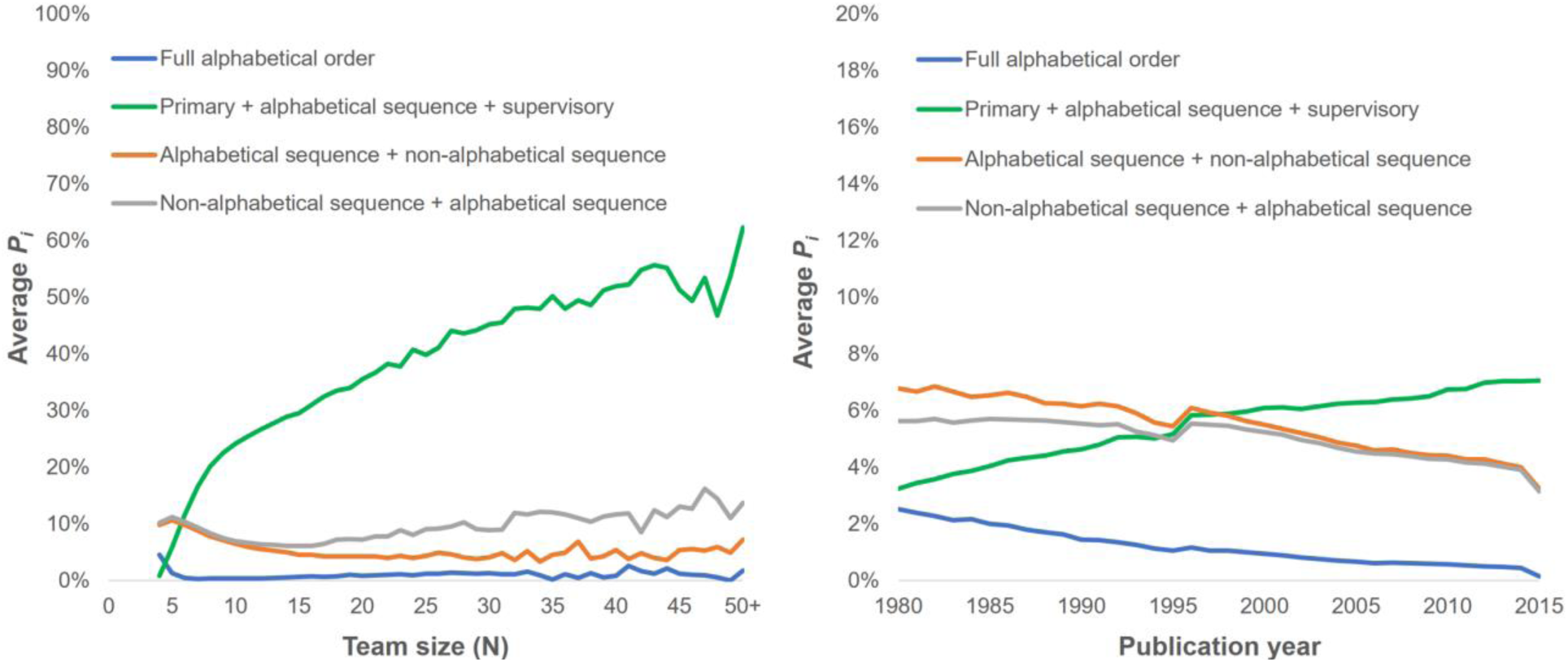
Proportion of article bylines containing intentional alphabetical order as a function of the number of authors (left), and publication year (right).

The right panel of indicates that the proportion of articles with alphabetically ordered middle authors has increased steadily over the last 35 years, rising from approximately 3% of articles in 1980 to almost 8% in 2015. Inversely, the number of bylines in full alphabetical order and of bylines where the first or last authors are in an alphabetical sequence have decreased.

The average size of teams producing biomedical articles varies over time, which may have an effect on the trends observed in Fig 3. To control for this variation, we performed a binomial logistic test to measure the effect of the team size and the publication year depending on the probability that the middle authors are ordered alphabetically on a byline with 95% certainty. In order to maintain this level of certainty, the test was performed on the subset of 2,527,997 articles authored by 7 to 100 individuals (the lowest *r* and *N* values for which *P_i_* ≥ .95 are 5 and 7, respectively). The model was statistically significant χ2(2) = 66,100.220, p < .001 but explained only 8.6% of the variance in the presence of an alphabetical sequence of authors in the bylines, correctly classifying only 3.4% of cases. As shown in table 1, the year of publication has in fact no effect on the proportion of bylines with alphabetically ordered middle authors (Exp(B) = 1.001). However, the team size does have an effect (Exp(B) = 1.145) and was the only statistically significant predictor, as shown in table 2.

**Table 1.** Logistic regression predicting the likelihood of an article containing alphabetically ordered middle authors based on the number of authors and the publication year.

|  | B | S.E. | Wald | df | Sig. | Exp(B) | 95% C.I. for EXP(B) |  |
| --- | --- | --- | --- | --- | --- | --- | --- | --- |
|  |  |  |  |  |  |  | Lower | Upper |
| Number of authors | .135 | .001 | 58258.286 | 1 | .000 | 1.145 | 1.143 | 1.146 |
| Publication year | .001 | .000 | 3.261 | 1 | .071 | 1.001 | 1.000 | 1.002 |
| Constant | -6.099 | .846 | 51.975 | 1 | .000 | .002 |  |  |

Having established the high prevalence of bylines containing alphabetically ordered middle authors, we proceeded to analyze the team composition of articles where such sequences are found, which limited our analysis to the 74,555 articles containing a single alphabetical sequence for which *P_i_* ≥ 0.95. Fig 4 displays average proportion of primary, middle and supervisory author as a function of the total team size (left) and of the publication year (right). Results suggest that independently of the team size, more than half the authors are middle authors. However, this proportion decreases slightly as the total team size increases. The other team members are distributed almost equally between primary and supervisory authors, the former being on average slightly more numerous than the former. Overall, the average team is composed of 20.9% (*SD* = .117) primary authors, 60.1% (*SD* = .141) middle authors, and 19.0% (*SD* = .119) supervisory authors.

**Fig 4.**
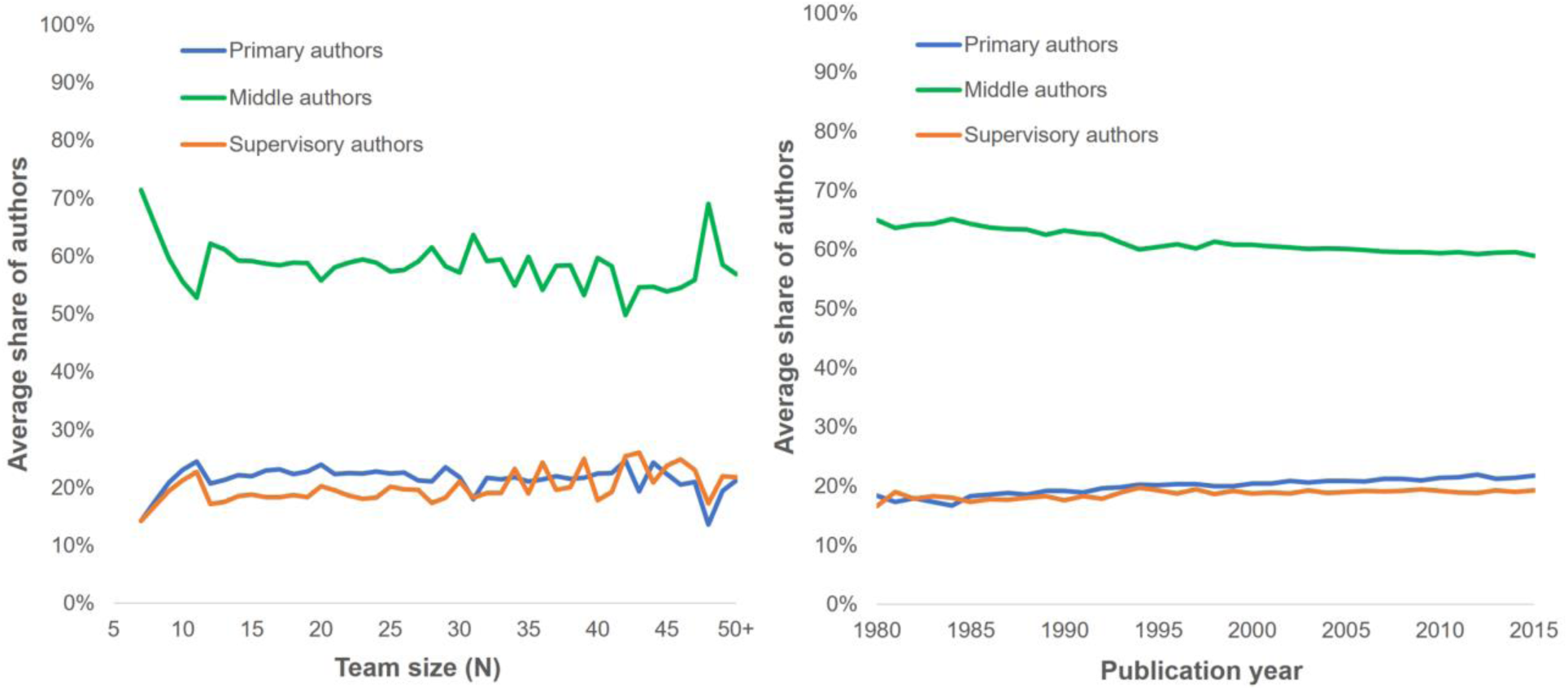
Average share of primary, middle and supervisory authors as a function of the total team size (left) and publication year (right).

The right panel of Fig 4 shows a slight decrease of the average proportion of middle authors in research teams, from 65.0% in 1980 to 59.0% in 2015. To disentangle the confounding effects of time and team size on the average proportion of middle authors, we performed a multiple regression to predict the share of middle authors from the total number of authors and the publication year. The model shows a low negative effect of both independent variables, with F(2, 105,530) = 2,584.516, p < .001, adj. R2 = .047. Regression coefficients and standard errors are shown in Table3.

**Table 2.** Coefficients of the multiple regression to predict the share of middle authors from the number of authors and the publication year.

|  | *B* | *SE<sub>B</sub>* | *β* | *P* |
| --- | --- | --- | --- | --- |
| Intercept | 4.458 | .125 |  |  |
| Number of authors | -.003 | .000 | -.185 | .000 |
| Publication year | -.002 | .000 | -.093 | .000 |

note: *B* = Unstandardized regression coefficient; *SE<sub>B</sub>* = Standard error of the coefficient; *β* = Standardized coefficient.

In this last part of our analysis, we look at how the overall contribution of middle authors to the biomedical literature has evolved over time. Past literature [e.g.,,3] has highlighted the growth of biomedical research teams. Our aim was to evaluate whether the relative number of middle authors have been increasing at a lower, similar or higher rate than the teams’ size. We calculate the middle authors’ contribution to biomedical research by dividing the sum of all *r* for which *P_i_* ≥ 0.95 by the sum of team size *N* for all articles published in a given year. Fig 5 shows a fourfold increase of the contribution of middle authors to the biomedical literature over the 1980–2015 period. While alphabetically ordered middle authors accounted for only 0.2% of all authors in 1980, they represented nearly 1.8% of authors in 2015, a ninefold increase. In comparison, the average team size has only doubled over the same period, going from 2.9 authors in 1980 to 6.6 authors in 2015. This suggests that over the last 35 years, middle authors have been playing an increasingly large part in the production of knowledge in the biomedical field.

**Fig 5.**
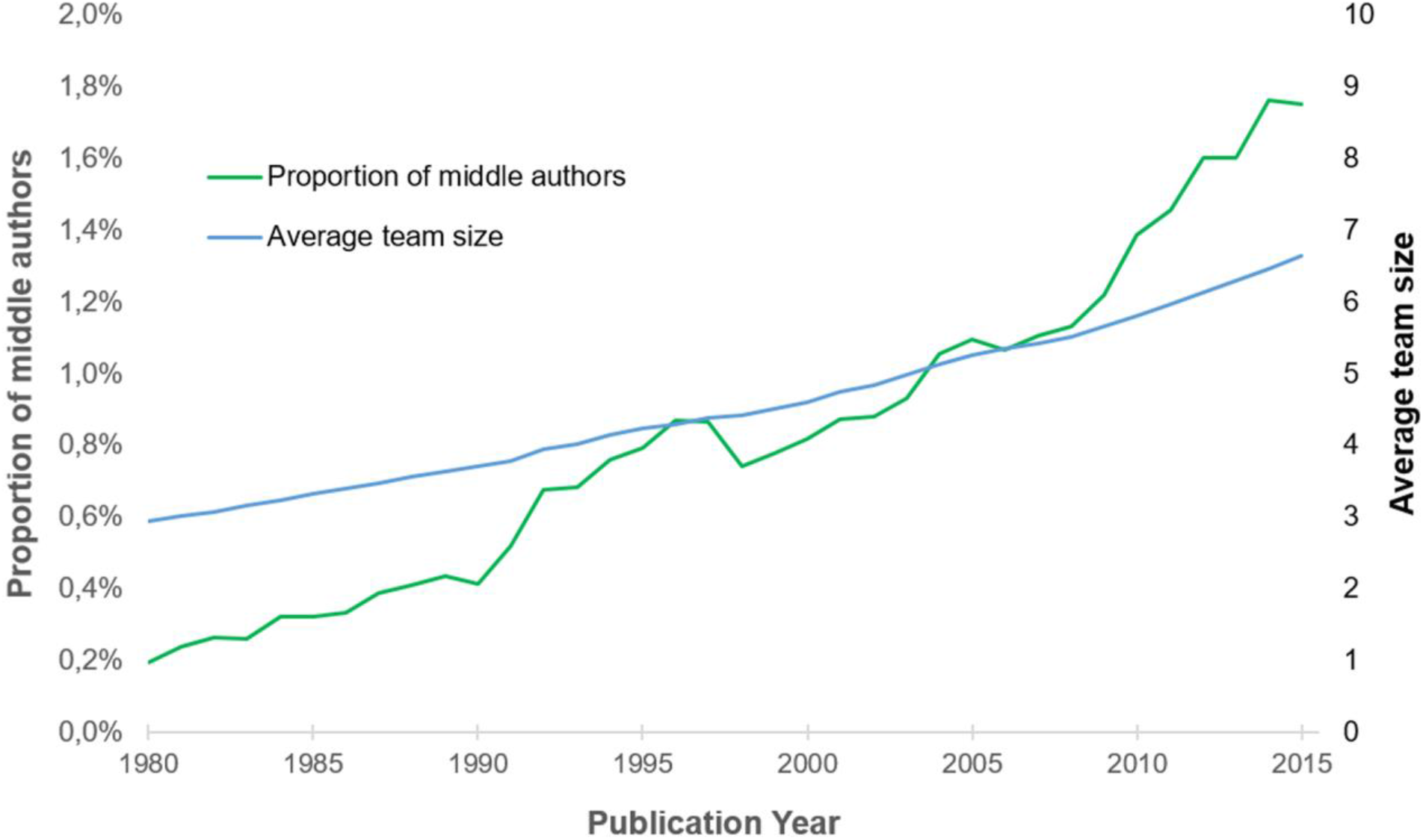
Overall relative contribution of middle authors to biomedical research.

## Discussion

The nature, scope and complexity of a research project determine the amount of work, the various tasks and the array of knowledge and skills required. These are determinant factors in establishing the size of the research team and the division of labor among its members. The naming and ordering of authors, which we use to assess the relative contribution of team members, is the product of a more or less normalized social process. Indeed, while strong (but implicit) disciplinary norms serve to guide authorship, other factors come to bear such as the existing relationships between collaborators and their position in the institutional hierarchy.

In the first part of our analysis, we show a clear relation between the size of teams and the prevalence of alphabetically ordered middle authors. This can be explained by two different factors. Firstly, all other things being equal, a larger team will lead to a greater division of the work. The usually uneven distribution of tasks will in turn determine author order, and allow for a distinction between primary, middle and supervisory authors. It is thus logical that the number of middle authors will increase as team increase in size. Secondly, since it is more difficult to order very large number of authors based on their contributions, especially when contributions are small and diverse, the use of alphabetical order to list most authors on large teams should logically prevail.

Results also show that, for papers with 7 authors or more, middle authors constitute the largest proportion of research teams. This supports the idea that work is unevenly distributed among team members: a few primary authors lead the experimental work, others have a supervisory role, and the rest of the authors share smaller and/or more technical parts of the work. However, a striking feature of the share of authors that fall in each of these three categories is their invariance as a function of team size, which suggests that growth in authors’ lists are not solely due to an increase in secondary contributions but, rather, in equivalent increases of all types of roles and contributions.

The large number of middle authors, especially in long authors’ list, also raises questions relating to authorship practices and criteria. It is possible, for instance, that the increasingly long bylines are not only reflecting an increase in collaboration, but also that small contributions are increasingly rewarded with authorship. Using ordering middle authors alphabetically may reduce the incentive to keep the author list as short as possible. Indeed, when authors are ordered according to their contribution or in full alphabetical order, adding a name on the byline reduces each authors’ share of credit. Clearly distinguishing primary, middle and supervisory authors by using alphabetical order does reduce this ‘loss of credit’ because primary and supervisory authors will remain differentiated from the middle authors, no matter the number of middle authors. Thus, listing middle authors in alphabetical order might increase the propensity to include more (middle) authors. Interestingly, this suggests that while the order of authors is determined by the type and amount of work, the method used for ordering names may also determine the type and amount of work required for an individual to be listed as an author. Another incentive for rewarding small contributions with authorship is the responsibility that is associated with authorship [17]. In a sense, naming all contributors as authors allows a more refined attribution of responsibility, where no author as to take responsibility for the work of others.

This idea of rewarding small (or all) contributions with authorship seems somewhat at odds with the current authorship guidelines of the International committee of Medical Journal Editors, which state that authorship is to be based on the following criteria:

- ‘Substantial contributions to the conception or design of the work; or the acquisition, analysis, or interpretation of data for the work; AND
- Drafting the work or revising it critically for important intellectual content; AND
- Final approval of the version to be published; AND
- Agreement to be accountable for all aspects of the work in ensuring that questions related to the accuracy or integrity of any part of the work are appropriately investigated and resolved.’ [28].

However, what constitutes a substantial contribution is left to the judgment of researchers. Also the guidelines do not take into account the different roles of individual team members, or the extent of their contribution. Nonetheless, as observed in our paper, authors *do* make this distinction, and it can be observed when they choose to list middle authors alphabetically. Furthermore, it seems plausible that in cases where very large numbers of authors are listed on a byline, and especially when middle authors are listed in alphabetical order, some authors will not meet the substantial contribution criterion. The scientific community’s apparent lack of adherence to authorship guidelines might suggest that it is time to further examine and ‘normalize’ emerging practice. It might indeed be more effective to promote an inclusive authorship model instead of the restrictive models that reward only main contributors and leave other contributions out of the byline.

## Conclusion

In this research, we demonstrated that the listing of middle authors alphabetically is a practice used frequently in biomedical research, especially for those articles with a large number of authors. We also showed that when middle authors are identified alphabetically, they represent on average more than half of the research team. This indicates that the author inflation might be due in part to increased division of labor, and not only to the growing complexity of research projects. This is reflected in the fact that the share of total authorships attributed to alphabetically ordered middle authors has increased more than average team size over the last 35 years. As discussed above, these results provide insights not only on collaboration and division of labor in biomedical research, but also on authorship practices. The increase in team size has raised issues that have been widely discussed, including the lack of transparency of authors’ contributions and the difficulties of assigning responsibility for the work as a whole or for its different parts. In addressing some of these issues, Baerlocher and colleagues [21] proposed that authors be designated as either primary, supervisory or contributing (middle) authors. We believe that such a system would be effective mainly in large teams as it would provide a normative framework that is more transparent and also better suited to reflect collaboration and division of labor in biomedical research. It would also recognize individuals who make smaller contributions as contributing authors; this inclusive approach is more representative of various contributions than the current ‘all or nothing’ model that excludes some contributors from the byline.

However, effective implementation of this model would involve its acceptance and adoption in research evaluation processes. Most of the currently used bibliometric indicators (e.g. the H-index) do not take into account one’s position on the byline. Consequently, being middle author may be paradoxically more rewarding, from a cost-benefit perspective, than being a primary or supervisory author. Inversely, indicators and evaluation processes that only put emphasis on the first position might create disputes and hinder collaboration and division of labor.

## S1. Supplement. Calculating the probability of chance and intentional alphabetical order

Let n ≥ 1 be an integer and let *Sn* denote the set of permutations of {1,…, *n*}. Let us write

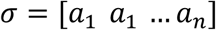

for the permutation with *σ(i)* = *a_i_*. For 1 ≤ *r* <= *n*, we shall say that σ has an increasing subsequence of length *r* if there exists *r* consecutive terms in increasing order:

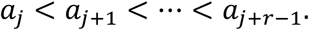

Let us denote 
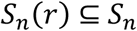
the subset consisting of those permutations with an increasing subset of length *r*. For instance, we have

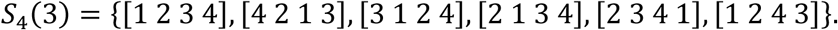

Let *s_n_*(r) denote the number of elements of *S_n_*(r), so for instance, we have *s*_4_(3) = 7. We have *s_n_*(r)= 1 since there can only be one entirely increasing permutation. On the other hand, we have *s_n_*(1)=!n because every permutation trivially contains a 1-term increasing subsequence.

Define 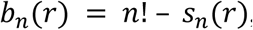
which is the number of permutations of *n* authors which do *not* contain an alphabetically ordered subsequence of *r* authors. One can show by a combinatorial argument (See Elizalde and Noy, 2003) that the exponential generating function

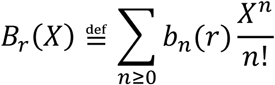

can be written as

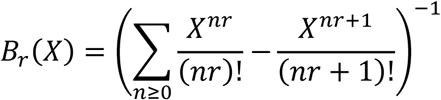

Using this formula, we computed *b_n_(r)* with the computer algebra system SAGE^1^. The Probability of intentional alphabetical order *Pi* is thus obtained with the following formula:

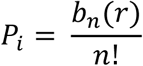

Inversely, the probability of chance alphabetical order *Pc* is obtained with the following formula:

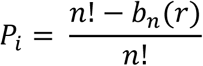

## Footnotes

1 Sage Mathematics Software (Version 7.1), The Sage Developers, http://www.sagemath.org.

## References

LeeS,BozemanB., The Impact of Research Collaboration on Scientific Productivity. Soc Stud Sci. 2005;35:673–702. doi:10.1177/0306312705052359.

Beaver D deB.,RosenR.Studies in scientific collaboration-Part II. Scientific co-authorship, research productivity and visibility in the French scientific elite, 1799–1830. Scientometrics. 1979;1: 133–149.

WuchtyS,JonesBF,UzziB., The Increasing Dominance of Teams in Production of Knowledge. Science. 2007;316: 1036–1039. doi:10.1126/science.1136099.

LarivièreV, GingrasY,SugimotoCR,TsouA.Team size matters: Collaboration and scientific impact since 1900. J Assoc Inf Sci Technol. 2015;66: 1323–1332. doi:10.1002/asi.23266.

JonesBF,WuchtyS,UzziB., Multi-University Research Teams: Shifting Impact, Geography, and Stratification in Science. Science. 2008;322: 1259–1262. doi:10.1126/science.1158357.

AdamsJD,BlackGC,ClemmonsJR,StephanPE. Scientific teams and institutional collaborations: Evidence from U.S. universities, 1981–1999. Res Policy. 2005;34: 259–285doi:10.1016/j.respol.2005.01.014.

DefazioD,LockettA,WrightM.Funding incentives, collaborative dynamics and scientific productivity: Evidence from the EU framework program. Res Policy. 2009;38: 293–305. doi:10.1016/j.respol.2008.11.008.

KatzJS,MartinBR., What is research collaboration? Res Policy. 1997;26: 1–18. doi:10.1016/S0048-7333(96)00917-1.

BozemanB,CorleyE.Scientists’ collaboration strategies: implications for scientific and technical human capital. Res Policy. 2004;33:599–616. doi:10.1016/j.respol.2004.01.008.

SmithD,KatzJS.,Collaborative Approaches to Research.A Report to the Higher Education Funding Council for England. 2000. Available: http://users.sussex.ac.uk/~sylvank/pubs/collc.pdf

CroninB.Hyperauthorship: A postmodern perversion or evidence of a structural shift in scholarly communication practices?J Am Soc Inf Sci Technol. 2001;52: 558–569. doi:10.1002/asi.1097.

MaienscheinJ.Why Collaborate ? J Hist Biol. Springer; 2012;26: 167–183.

SmithCG. Scientific Performance and the Composition of Research Teams. Adm Sci Q. 1971;16: 486. doi:10.2307/2391768.

LarivièreV,DesrochersN,MacalusoB, MongeonP,Paul-HusA,SugimotoCR.Contributorship and division of labor in knowledge production. Soc Stud Sci. 2016;46: 417–435. doi:10.1177/0306312716650046.

RennieD,YankV.If authors became contributors, everyone would gain, especially the reader. Am J Public Health. American Public Health Association; 1998;88: 828–830.

CroninB.Rates of return to citation. J Doc. 1996;52: 188–197. doi:10.1108/eb026967.

BirnholtzJP. What does it mean to be an author? The intersection of credit, contribution, and collaboration in science. J Am Soc Inf Sci Technol. 2006;57: 1758–1770. doi:10.1002/asi.20380.

MertonRK. Priorities in scientific discovery: a chapter in the sociology of science. Am Sociol Rev. 1957;22: 635–659.

Knorr-CetinaK.Epistemic cultures: Forms of reason in science. Hist Polit Econ. 1991;23: 105–122.

PontilleD.La signature scientifique : une sociologie pragmatique de l’attribution. Paris: CNRS; 2004.

BaerlocherMO,NewtonM, GautamT, TomlinsonG, DetskyAS., The Meaning of Author Order in Medical Research. J Investig Med. 2007;55: 174–180. doi:10.2310/6650.2007.06044

MongeonP,LarivièreV.Costly collaborations: The impact of scientific fraud on co-authors’ careers. J Assoc Inf Sci Technol. 2016;67: 535–542. doi:10.1002/asi.23421.

BoyackKW,KlavansR,SorensenAA,IoannidisJPA. A list of highly influential biomedical researchers, 1996-2011. Eur J Clin Invest. 2013;43: 1339–1365. doi:10.1111/eci.12171.

WarnerET,CarapinhaR,WeberGM,HillE V.,ReedeJY., Faculty Promotion and Attrition: The Importance of Coauthor Network Reach at an Academic Medical Center. J Gen Intern Med. 2016;31: 60–67. doi:10.1007/s11606-015-3463-7.

PontilleD. Qu’est-ce qu’un auteur scientifique. Sci la Société. 2006;67: 77–93.

ZuckermanHA. Patterns of Name Ordering Among Authors of Scientific Papers: A Study of Social Symbolism and Its Ambiguity. Am J Sociol. The University of Chicago Press; 1968;74: 276–291. doi:10.2307/2775535.

WaltmanL.An empirical analysis of the use of alphabetical authorship in scientific publishing. J Informetr. 2012;6: 700–711. doi:10.1016/j.joi.2012.07.008.

International Committee of Medical Journal Editors. Recommendations for the conduct, reporting, editing, and publication of scholarly work in medical journals [Internet]. 2015 [cited 28 Aug 2016]. Available:http://www.icmje.org/icmje-recommendations.pdf.

Elizalde, S.,Noy, M. (2003). Consecutive patterns in permutations. Advances in Applied Mathematics, 30(1–2), 110–125.

